# Genome-wide prediction of topoisomerase II*β* binding by architectural factors and chromatin accessibility

**DOI:** 10.1101/2020.03.23.003277

**Authors:** Pedro Manuel Martínez-García, Miguel García-Torres, Federico Divina, José Terrón-Bautista, Irene Delgado-Sainz, Francisco Gómez-Vela, Felipe Cortés-Ledesma

## Abstract

DNA topoisomerase II-*β* (TOP2B) is fundamental to remove topological problems linked to DNA metabolism and 3D chromatin architecture, but its cut-and-reseal catalytic mechanism can accidentally cause DNA double-strand breaks (DSBs) that can seriously compromise genome integrity. Understanding the factors that determine the genome-wide distribution of TOP2B is therefore not only essential for a complete knowledge of genome dynamics and organization, but also for the implications of TOP2-induced DSBs in the origin of oncogenic translocations and other types of chromosomal rearrangements. Here, we conduct a machine-learning approach for the prediction of TOP2B binding sites using publicly available sequencing data. We achieve highly accurate predictions, with accessible chromatin and architectural factors being the most informative features. Strikingly, TOP2B is sufficiently explained by only three features: DNase I hypersensitivity, CTCF and cohesin binding, for which genome-wide data are widely available. Based on this, we develop a predictive model for TOP2B genome-wide binding that can be used across cell lines and species, and generate virtual probability tracks that accurately mirror experimental ChIP-seq data. Our results deepen our knowledge on how the accessibility and 3D organization of chromatin determine TOP2B function, and constitute a proof of principle regarding the *in silico* prediction of sequence-independent chromatin-binding factors.

**Author summary:** Type II DNA topoisomerases (TOP2) are a double-edged sword. They solve topological problems in the form of supercoiling, knots and tangles that inevitably accompany genome metabolism, but they do so at the cost of transiently cleaving DNA, with the risk that this entails for genome integrity, and the serious consequences for human health, such as neurodegeneration, developmental disorders or predisposition to cancer. A comprehensive analysis of TOP2 distribution throughout the genome is therefore essential for a deep understanding of its function and regulation, and how this can affect genome dynamics and stability. Here, we use machine learning to thoroughly explore genome-wide binding of TOP2B, a vertebrate TOP2 paralog that has been linked to genome organization and cancer-associated translocations. Our analysis shows that TOP2B-DNA binding can be accurately predicted exclusively using information on DNA accessibility and binding of genome-architecture factors. We show that such information is enough to generate virtual maps of TOP2B binding along the genome, which we validate with *de novo* experimental data. Our results highlight the importance of TOP2B for accessibility and 3D organization of chromatin, and show that computationally predicted TOP2 maps can be accurately obtained using minimal publicly available datasets, opening the door for their use in different organisms, cell types and conditions with experimental and/or clinical relevance.

## Introduction

Type II DNA topoisomerases are unique in their ability to catalyze duplex DNA passage, and therefore the only enzymes capable of dealing with superhelical DNA structures, as well as unknotting and decatenating DNA molecules [1]. This places them in a privileged position to integrate and coordinate many aspects of genome organization and metabolism, and have indeed been related to virtually all aspects of chromatin dynamics [2, 3]. While the genomes of invertebrates and lower eukaryotic systems code for only one type II topoisomerase (TOP2), vertebrates encode two close paralogs (TOP2A and TOP2B) with similar catalytic and structural properties but with different roles [2, 4]. TOP2A is essential for cellular viability [5] and is mainly expressed in dividing cells [6], where it is required for chromosome segregation [2].

TOP2B, in contrast, is not essential at a cellular level [7], but is ubiquitously expressed and required for organismal viability due to roles in the transcriptional regulation of genes required for proper development of the nervous system [2, 6, 8]. Consistent with regulatory functions in transcription, TOP2B significantly associates with promoters and DNase I hypersensitivity sites, as well actively regulated epigenetic modifications and enhancers [9–13]. Furthermore, recent studies have linked TOP2B with 3D genome architecture due to a strong colocalization with architectural factors such as CTCF and the cohesin complex protein RAD21 at the boundaries of topologically associated domains (TADs), which has been interpreted as TOP2B functions in resolving topological problems derived from the active organization of the genome in the context of the loop extrusion model [11, 12].

Despite these fundamental roles in the organization and dynamics of the genome, TOP2 function can also pose a threat to its integrity [4]. Aberrant activity of TOP2, which can occur spontaneously or as a consequence of cancer chemotherapy, results in the formation of DNA double-strand breaks (DSBs). These are highly cytotoxic lesions that, if inefficiently or aberrantly repaired, can seriously compromise cell survival and genome stability. Indeed, unrepaired or misrepaired DSBs can lead to genome rearrangements associated with neurodegeneration, sterility, developmental disorders and predisposition to cancer [14–16]. In particular, transcription-associated TOP2B function at TAD boundaries has been recently linked to oncogenic chromosomal translocations [17, 18]. The specific functions of TOP2B in these processes are, however, still largely unexplored and far from being fully understood. A deep understanding of TOP2B function and regulation in a genome-wide context, and how it can result in chromosome fragility and alterations, will require thorough and unbiased studies, integrating different systems, cellular types and conditions.

In this work, we take advantage of the wealth of high-throughput sequencing data to comprehensively study the predictive power of a wide set of chromatin features to identify TOP2B binding sites. We show that TOP2B localization can be accurately predicted using publicly available sequencing data, and identify architectural and open chromatin factors as the most informative features. Moreover, feature selection analysis indicates that TOP2B can be faithfully predicted using only three sequencing datasets. Indeed, models trained with DNase-seq and ChIP-seq of CTCF and RAD21 allow for accurate prediction across different mouse cell types, supporting a generalizing association of TOP2B with these features. Finally, we perform ChIP-seq of TOP2B in mouse thymocytes and human MCF7 and generate predictions on such cell types in order to validate our model. Genome wide predictive tracks accurately mirror experimental TOP2B tracks in both organisms, showing that our model can be used to generate virtual TOP2B signals in potentially any mammalian cell type and condition for which sequencing data of CTCF, RAD21 and DNase I hypersensitivity are available.

## Materials and methods

### Processing publicly available data

The experimental data used in this study is summarized in S1 Table. When available, we batch-downloaded mm9 and hg19 BAM files from ENCODE. Raw sequencing data was processed by first quality filtering and merging the reads of biological replicates and corresponding input samples (in the case of ChIP-seq experiments). Then we mapped them to the mouse or human genome (mm9 or hg19) using Bowtie 1.2 [19] with option “-m 1” so that reads that mapped only once to the genome were retained. Genome track plots of Fig 1 and S1 Fig were generated using Bioconductor packages Gviz [20] and trackViewer [21].

**Fig 1.**
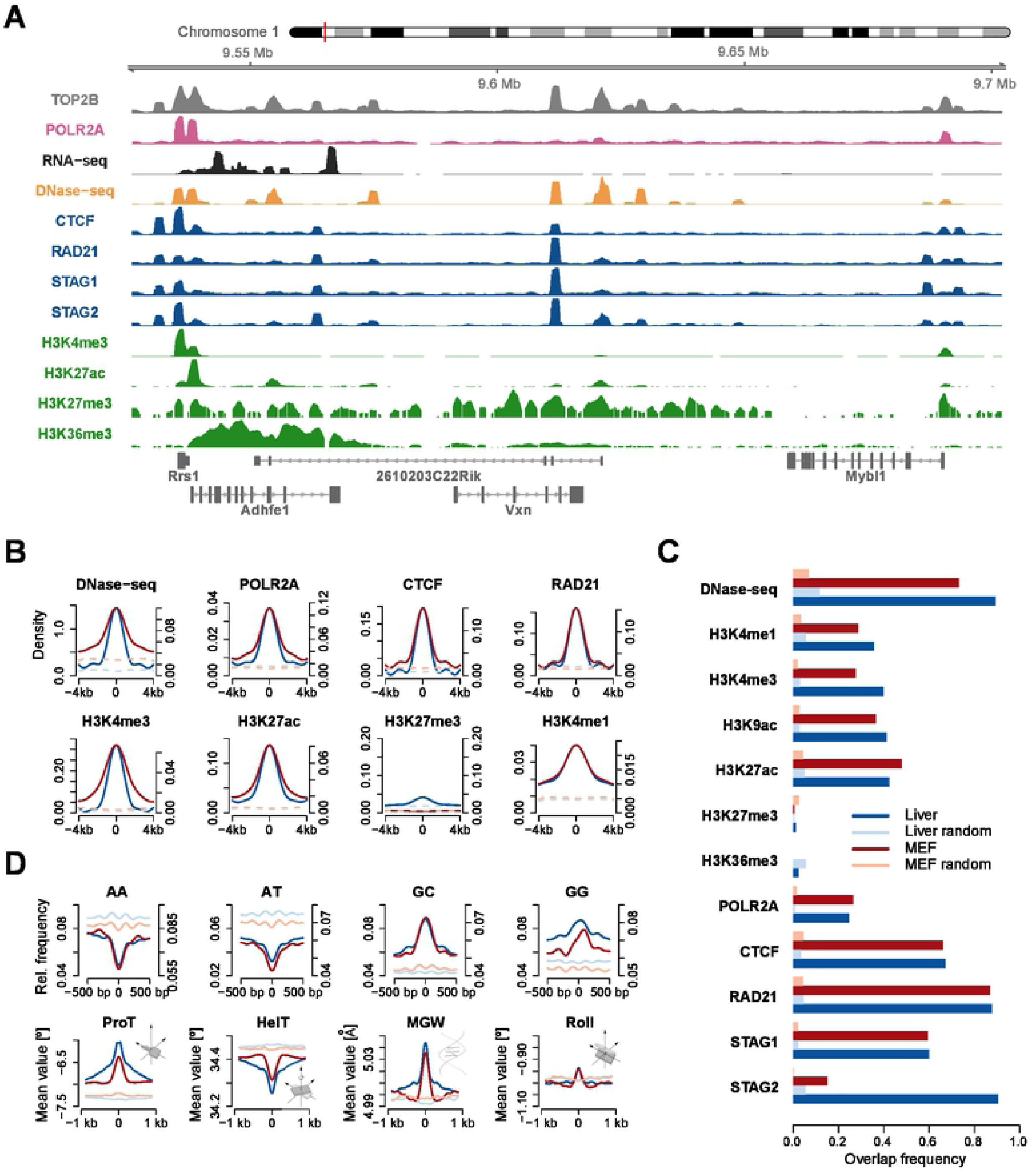
Chromatin landscape of TOP2B. **A**. Genome browser view for TOP2B ChIP-seq signal and other relevant chromatin features in mouse liver (http://genome.ucsc.edu). **B**. Average reads enrichment of selected chromatin features within *±*4 kb of TOP2B binding center (solid lines) and random regions (dashed lines) in mouse liver (blue) and MEFs (red). Signals were smoothed using nucleR [39]. Left and right sided ordinate axes correspond to mouse liver and MEFs, respectively. **C**. Overlap frequencies of chromatin features at TOP2B binding sites as compared to random regions in mouse liver and MEFs. **D**. Top panel shows relative frequency of several polynucleotides and dinucleotides within *±*500 bp of TOP2B binding center and random regions. Bottom panel shows average of 3D DNA shape features within ±1 kb of TOP2B binding center and random regions. Profiles corresponding to MEFs and liver are colored as in C.

### Identification of TOP2B Binding Sites

We used HOMER [22] for the computation of TOP2B peaks and retained those with a fold change over the input sample greater than 50. Then, peaks overlapping *>* 20% of their width with non-mappable regions or regions of the ENCODE blacklist [23] were discarded. When replicates were available, peaks were called for individual replicates and overlapping peaks were kept. For each training cell type, we generated the same number of random regions in the genome than TOP2B binding sites were identified. Such regions had 300 bp length and were selected so that they had the same distribution across chromosomes as TOP2B peaks. Again, non-mappable regions and regions of the ENCODE blacklist were discarded. To make sure they did not overlap potential TOP2B peaks that were not detected by the peak calling algorithms, regions with high levels of TOP2B/INPUT ChIP-seq signal were also discarded. In order to account for sequence composition biases, an additional set of random regions was generated so that their distribution of sequence G+C content matched that of TOP2B peaks.

### Quantification of High-Throughput Sequencing Data

To score sequencing data around TOP2B binding sites, we based on the scoring approach of [24]. For ChIP-seq experiments, reads aligned to a 300 bp window centered on TOP2B binding center were first considered, normalized to the size of the corresponding sequencing library and scaled to the size of the smaller library (among test and input). Then, they were added 1 as a pseudocount (to avoid undefined values in posterior logarithmic transformations) and the ratio of test versus input signal was calculated. Finally, this value was log transformed. RNA-seq, DNase-seq and MNase-seq were scored as the number of reads aligned within the same 300 bp window divided by the library size. CpG methylation was measured as the total percentage of methylated CG for each CpG dinucleotide as described in [25].

### DNA shape and sequence features

Three-dimensional DNA shape features at base pair or base pair step resolution were derived from the high-throughput approach implemented in the Bioconductor library DNAshapeR [26]. We used thirteen DNA shape features: helix twist, propeller twist, minor groove width, roll, shift, slide, rise, tilt, shear, stretch, stagger, buckle and opening [27]. For a given DNA sequence of length L, the k-mer feature at each nucleotide position was encoded as a binary vector of length 4^*k*^, as described in [28]. A value of 1 represents the occurrence of a particular k-mer from starting at that position. The k-mer features for a whole DNA sequence were then encoded by the concatenation of the k-mer feature vectors at each position of the sequence. As a result, we obtained a binary vector of length 4^*k*^(L-k+1).

### Machine learning

#### Training data matrix

After scoring the sequencing experiments as well as DNA shape and sequence parameters, we built up a final data matrix with rows representing TOP2B/random sites and columns corresponding to scored features. Since we identified a total of 13, 128 and 8, 413 TOP2B peaks in mouse liver and MEFs respectively, we ended up with model matrices of 32, 766 columns and either 26, 256 (mouse liver) or 16, 826 (MEFs) rows.

#### Classifiers

The Naive Bayes [29] (NB) classifier applies the Bayes rule to compute the probability of a class label given an instance. For such purpose, the classification model learns, from training data, the conditional probability of each feature given the class label. NB assumes that all features are conditionally independent given the values of the class.

Support Vector Machine [30] (SVM) is a classifier based on the idea of finding a hyperplane that optimally separates the data into two categories by solving an optimization problem. Such hyperplane, is the one that maximizes the margin between classes. In case of non-separable data, it optimizes a weighted combination of the misclassification rate and the distance of the decision boundary to any sample vector. After several tests with different kernels (linear, polynomial, radial basis and sigmoid), we use a linear kernel due to its good performance and simplicity.

Random Forests [31] is a widely applied tree-based supervised machine-learning algorithm designed as a combination of bagging and a random selection of features that has proven to be an effective tool for the statistical analysis of high-dimensional genomic data [32].

#### Feature selection algorithms

Fast Correlation Based Filter [33] (FCBF) is an efficient heuristic that, based on information theory measures, performs a relevance and redundancy analysis for selecting a good subset of features. Basically, FCBF is a backward search strategy that uses Symmetrical Uncertainty (SU) as goodness function to measure non-linear dependencies between features. In order to identify relevant features, it requires to set a threshold parameter *δ* so that only values above *δ* are considered relevant.In this work we set *δ* = 0 since there is no rule about this parameter tuning and in the datasets under study only a small subset of features have a SU value different to 0.

Scatter Search [34, 35] (SS) is a population-based metaheuristic that uses intensification and diversification mechanisms to generate new solutions. The strategy starts by generating an initial population of diverse solutions from the solution space. Then, it selects a subset of features, called Reference Set (RefSet), of high quality and dispersed solutions from the initial population. In the next step the method selects all subsets consisting of two solutions and combine them in order to generate new solutions that are improved by applying a local search which will yield to local optima. Finally, a static update of the reference set is carried out selecting solutions according to quality and diversity from the union of the original RefSet and the new local optima found.

In this work we use the Correlation Feature Selection [36] (CFS) measure as goodness function. CFS evaluates subsets of features by considering the non-linear correlation Symmetrical Uncertainty function.

### Genome wide predictions

Once we identified the most predictive features, a generalized TOP2B binding model was built by training on DNase-seq, RAD21 and CTCF data from mouse liver, MEFs and activated B cells using Random Forests [31]. We first used 5-fold cross-validation to confirm that the model was able to accurately predict TOP2B in the training cell types (as shown in Results and Discussion; Table 4) and then applied the model to the mouse thymus and the human MCF7 mappable genomes. First, we splitted the genomes into bins of 300 bp with sliding windows of 50 bp. Then we scanned each bin with our model and obtained a TOP2B binding probability vector that was used to build up a bedgraph file. Probability track plots of Fig 6 and S6-7 Figures were generated using UCSC genome browser [37].

### ChIP-seq

Cells were fixed by adding formaldehyde to a final concentration of 1% and incubated at 37uC for 10 min. Fixation was quenched by adding glycine to a final concentration of 125 mM at cell culture plates. After two washes with cold PBS, in the presence of complete protease inhibitor cocktail (Roche) and PMSF, cell pellet was lysed in two steps using 0.5% NP-40 buffer for nucleus isolation and SDS 1% lysis buffer for nuclear lysis. Sonication was performed using Bioruptor (Diagenode, UCD-200) at high intensity and three cycles of 10 min (30” sonication, 30” pause) and chromatin was clarified by centrifugation (17000xg, 10 min, 4 ºC). For IP, 30 µg chromatin and 4 µg antibody (anti-TOP2B, SIGMA) were incubated o/n in IP buffer (0.1% SDS, 1% TX-100, 2 mM EDTA, 20 mM TrisHCl pH8, 150 mM NaCl) at 4ºC, and then with 25 µL of pre-blocked (1 mg/ml BSA) Dynabeads protein A and Dynabeads protein G (ThermoFisher) for 4 hours. Beads were then sequentially washed with IP buffer, IP buffer containing 500 mM NaCl and LiCl buffer (0.25 M LiCL, 1% NP40, 1% NaDoc, 20 mM TrisHCl pH8 and 1 mM EDTA). ChIPmentation was carried out as previously described (Schmidl 2015) using Tagment DNA Enzyme provided by the Proteomic Service of CABD (Centro Andaluz de Biología del Desarrollo). DNA was eluted by incubation at 50ºC in 100 *µ*L elution buffer (1% SDS, 100 mM NaHCO3) for 30 min followed by an incubation with 200 mM NaCl and 10 µg of Proteinase K (ThermoFisher) to revert cross-linking. DNA was then purified using Qiagen PCR Purification columns. Libraries were amplified for N-1 cycles (being N the optimum Cq determined by qPCR reaction) using NEBNext High-Fidelity Polymerase (M0541, New England Biolabs), purified and size-selected using Sera-Mag Select (GE Healthcare) and sequenced using Illumina NextSeq 500 in a single-end configuration. ChIP-seq datasets are available under GEO accession number GSE141528.

## Results and Discussion

### Chromatin landscape of TOP2B binding sites

We started by comprehensively assessing the chromatin landscape of TOP2B binding sites in two mouse cell types: embryonic fibroblasts (MEFs) and liver. We collected high-throughput sequencing data from ENCODE [38] and several independent studies (S1 Table) and observed the expected colocalization with open chromatin sites, RNA polymerase II (Pol2), architectural components and transcription associated histone modifications; genome tracks in both cell types are provided as an example (Fig 1A; S1 Fig). Then, we studied the position specific distribution of each chromatin feature around the TOP2B binding center (Fig 1B). Several marks were found to be highly enriched in both cell types, such as DNase-seq, CTCF and RAD21. They also showed a high colocalization frequency with TOP2B (Fig 1C). For instance, 89% of TOP2B sites in mouse liver colocalized with DNase-seq peaks, while for randomly selected regions this percentage decreased to 11%. Similarly, 87% of TOP2B peaks overlapped with RAD21 in MEFs as compared to 4% of random regions. These observations are in line with previous findings reporting a preferential association of TOP2B with accessible chromatin, actively transcribed promoters and regulators of genome architecture [10, 11], and suggest that chromatin features measured by next generation sequencing provide valuable information for the prediction of TOP2B binding.

Current experimental data suggest that TOP2B does not bind to a specific DNA motif [11], which agrees with topoisomerases specifically acting where DNA topological problems arise. However, TOP2B associates with DNA regions that are frequently bound by sequence-specific transcription factors [9–11], so the fact that particular topological structures can be favored by specific properties of DNA sequence is still a possibility. Indeed, a recent study has provided evidence that DNA sequence governs the location of supercoiled DNA [40]. To explore whether sequence composition of TOP2B binding sites can be used in our predictive approach, we analyzed the spatial distribution of DNA sequence dinucleotides within 1kb windows centered on TOP2B peaks. For both murine cell types, TOP2B sites showed an increase of GC and GG dinucleotides while AT and AA were depleted (Fig 1D, top). This result agrees with previously reported GC-rich motifs, such as CTCF and ESR1, at TOP2B occupied regions [10, 11].

Another feature that could potentially help in the prediction of TOP2B binding is DNA shape, which has been shown to affect the binding preferences of a number of DNA associated proteins [28, 41, 42]. In this sense, models trained with information on DNA shape have demonstrated significant improvements in transcription factor binding predictions [28]. To further inspect whether the observed sequence preferences of TOP2B sites are accompanied by specific 3D conformational parameters, we derived high-throughput predictions of DNA shape features from TOP2B peaks. Position specific profiles of such features revealed a specific pattern around the TOP2B binding center, characterized by decreased helix twist, increased minor groove width and propeller twist, and a modest enrichment of roll (Fig 1D, bottom). Such conformation suggests helical unwinding as consequence of a decrease in helix twist, which together with the widening of the minor groove could be providing an energetically favorable scenario for TOP2B binding.

Finally, we also interrogated whether CpG methylation is informative for TOP2B binding prediction. CpG methylation is a well-known epigenetic mark that has proven predictive ability for the localization of DNA binding proteins [43–45]. We profiled whole genome bisulfite sequencing (WGBS) data around TOP2B binding sites and found decreased CpG methylation as compared to random regions (S2 Fig), showing that this feature is likely to be informative for TOP2B prediction.

### A predictive model for TOP2B binding

In order to assess whether the above chromatin features could discriminate TOP2B binding sites from the rest of the genome, we applied the computational approach outlined in Fig 2. For both MEFs and liver, we first identified TOP2B peaks, resized them to 300 bp and generated the same number of random genomic regions (see Materials and methods). Then we scored 15 high-throughput sequencing experiments (S1 Table) together with DNA sequence and shape features within such regions. This resulted in a data matrix with rows representing TOP2B/random sites and columns corresponding to scored features. Finally, binary classifiers were trained and tested using 5-fold cross-validation.

**Fig 2.**
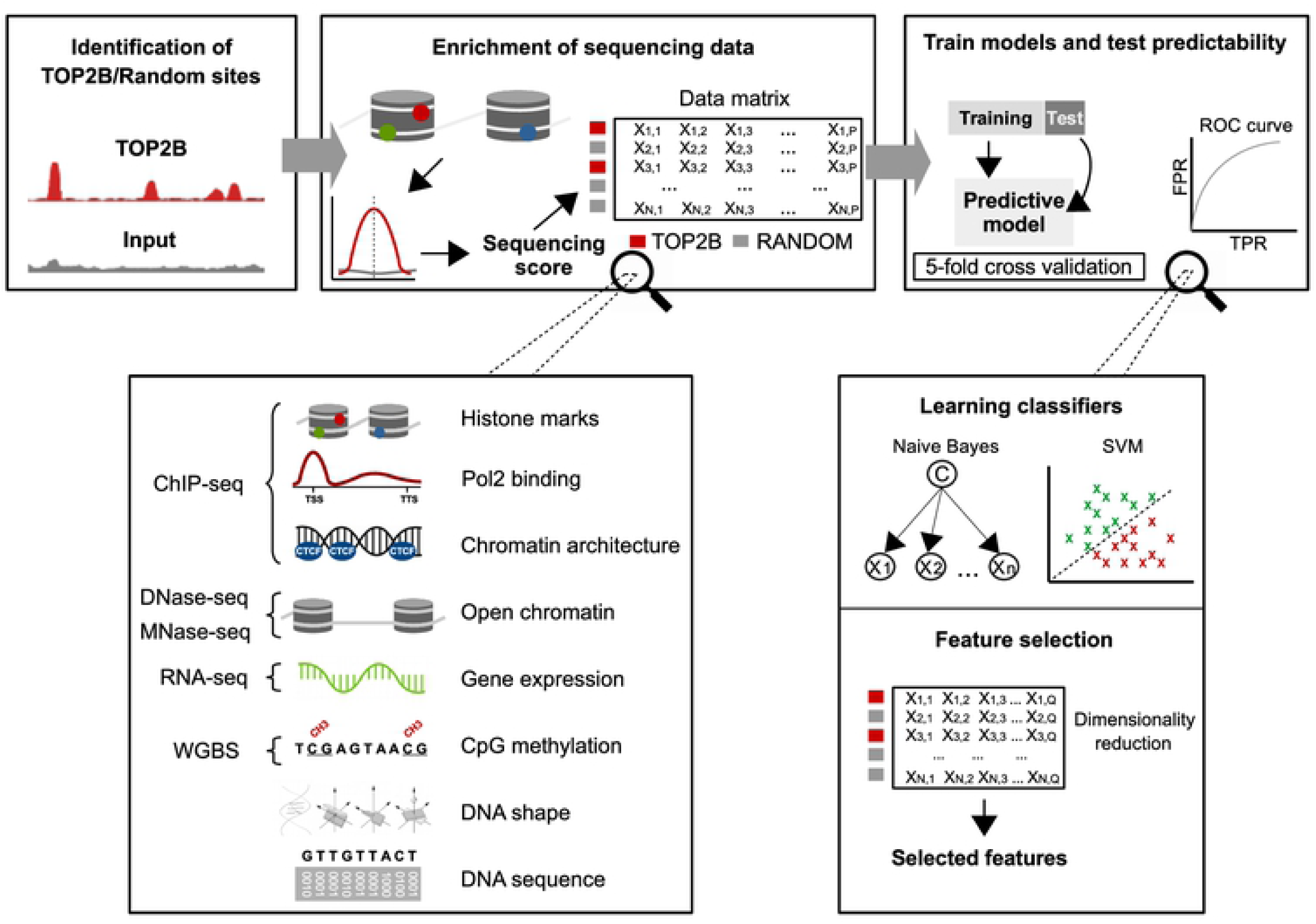
Machine learning schema for the prediction of TOP2B binding. TOP2B binding sites and random regions were first identified. Then, 15 high-throughput sequencing experiments together with DNA sequence and shape features were scored around such regions, which resulted a data matrix with rows representing TOP2B/random sites and columns representing the scored features. Finally, binary classifiers were trained and tested using 5 fold cross-validation and feature selection was applied to identify the most informative features.

In order to evaluate if DNA sequence is informative for the prediction of TOP2B binding, we represented DNA 1-mers, 2-mers and 3-mers for each nucleotide position in the TOP2B binding sites as described in [28] (see Materials and methods). This produced 1, 200 parameters representing 1-mers, 3, 584 parameters representing 2-mers and 13, 088 parameters representing 3-mers. We also included information on 13 DNA shape features using DNAshape method [46], which added other 7, 695 parameters to the model. This parametrization allowed us to measure the predictive ability of DNA sequence and shape to explain TOP2B binding with an unprecedented resolution. As a result, we ended up with model matrices of 32, 766 columns and either 13, 128 (liver) or 8, 413 (MEFs) rows.

### Chromatin features predict TOP2B

Since TOP2B binding regions display increased G+C content as compared to the average of the mouse genome, we trained one more model in which random regions were selected so that their distribution of sequence G+C content matches that of TOP2B peaks. This allowed us to account for potential biases of the predictions due to differences in G+C content. We employed Support Vector Machine (SVM) [30] and Naive Bayes (NB) [29] to build up the predictive models and used 5-fold cross-validation to estimate the prediction accuracy. We started by training on the whole set of features. Accurate predictions were obtained for the two murine cell types using both the GC-corrected and the regular model (Tables 1 and 2; Fig 3; S3 Fig), indicating that chromatin features can faithfully explain TOP2B binding. As a matter of fact, SVM trained on all features performed considerably better than NB. For instance, while regular models trained with NB achieved accuracies of 71% and 77.4% for MEF and liver, respectively, these values increased to 98% and 97.4% when using SVM. Similarly, accuracies obtained training the GC-corrected models with NB were 74.1% and 76.2% for MEF and liver, respectively, whereas SVM produced accuracies of 96.7% and 93%. Such performance differences are likely to be explained by many features showing dependencies to each other or being poorly informative. This would impair NB performance, which assumes all features are independent. Indeed, removal of not informative features leads to a significant increase of accuracies achieved by the NB algorithm (see next sections).

**Table 1.** Performance of SVM and NB classifiers in MEF.

| Cell type | Category | GC corrected |  | Random |  |
| --- | --- | --- | --- | --- | --- |
|  |  | NB | SVM | NB | SVM |
| MEF | All | 74.18 ± 1.08 | 96.72 ± 0.18 | 70.94 ± 0.75 | 98.06 ± 0.28 |
|  | Sequence | 59.15 ± 0.61 | 55.44 ± 1.10 | 66.87 ± 0.88 | 64.41 ± 0.48 |
|  | 3D shape | 55.08 ± 0.92 | 54.32 ± 0.87 | 66.23 ± 0.54 | 61.80 ± 0.97 |
|  | Histone marks | 84.82 ± 0.73 | 83.90 ± 0.66 | 88.61 ± 0.70 | 87.57 ± 0.74 |
|  | Architecture | 90.35 ± 0.61 | 94.98 ± 0.32 | 93.15 ± 0.58 | 96.52 ± 0.39 |
|  | Open chromatin | 81.92 ± 0.58 | 83.96 ± 1.44 | 90.48 ± 0.61 | 92.35 ± 0.63 |
|  | Expression | 50.36 ± 0.25 | 50.48 ± 0.54 | 50.77 ± 0.22 | 52.52 ± 1.57 |
|  | Pol2 | 60.60 ± 0.62 | 61.48 ± 1.06 | 57.71 ± 0.46 | 56.09 ± 0.92 |
|  | CpG methylation | 69.45 ± 0.88 | 73.82 ± 0.88 | 63.98 ± 0.79 | 68.90 ± 1.09 |

**Table 2.** Performance of SVM and NB classifiers in mouse liver.

| Cell type | Category | GC corrected |  | Random |  |
| --- | --- | --- | --- | --- | --- |
|  |  | NB | SVM | NB | SVM |
| Liver | All | 76.20 ± 0.97 | 93.00 ± 0.56 | 77.43 ± 0.20 | 97.47 ± 0.13 |
|  | Sequence | 62.95 ± 0.70 | 56.64 ± 1.10 | 74.98 ± 0.31 | 72.78 ± 0.94 |
|  | 3D shape | 57.97 ± 0.66 | 54.00 ± 1.42 | 73.19 ± 0.38 | 69.71 ± 0.46 |
|  | Histone marks | 78.03 ± 0.32 | 81.17 ± 0.47 | 78.97 ± 0.44 | 85.34 ± 0.41 |
|  | Architecture | 87.45 ± 0.53 | 91.74 ± 0.37 | 89.50 ± 0.54 | 94.26 ± 0.56 |
|  | Open chromatin | 71.68 ± 0.46 | 85.32 ± 4.22 | 83.35 ± 0.92 | 92.01 ± 1.57 |
|  | Expression | 50.00 ± 0.01 | 50.00 ± 0.01 | 50.04 ± 0.28 | 50.02 ± 0.01 |
|  | Pol2 | 54.92 ± 0.24 | 53.97 ± 0.42 | 52.95 ± 0.73 | 52.63 ± 0.90 |
|  | CpG methylation | 71.58 ± 0.34 | 75.31 ± 0.34 | 66.56 ± 0.36 | 69.58 ± 0.44 |

**Fig 3.**
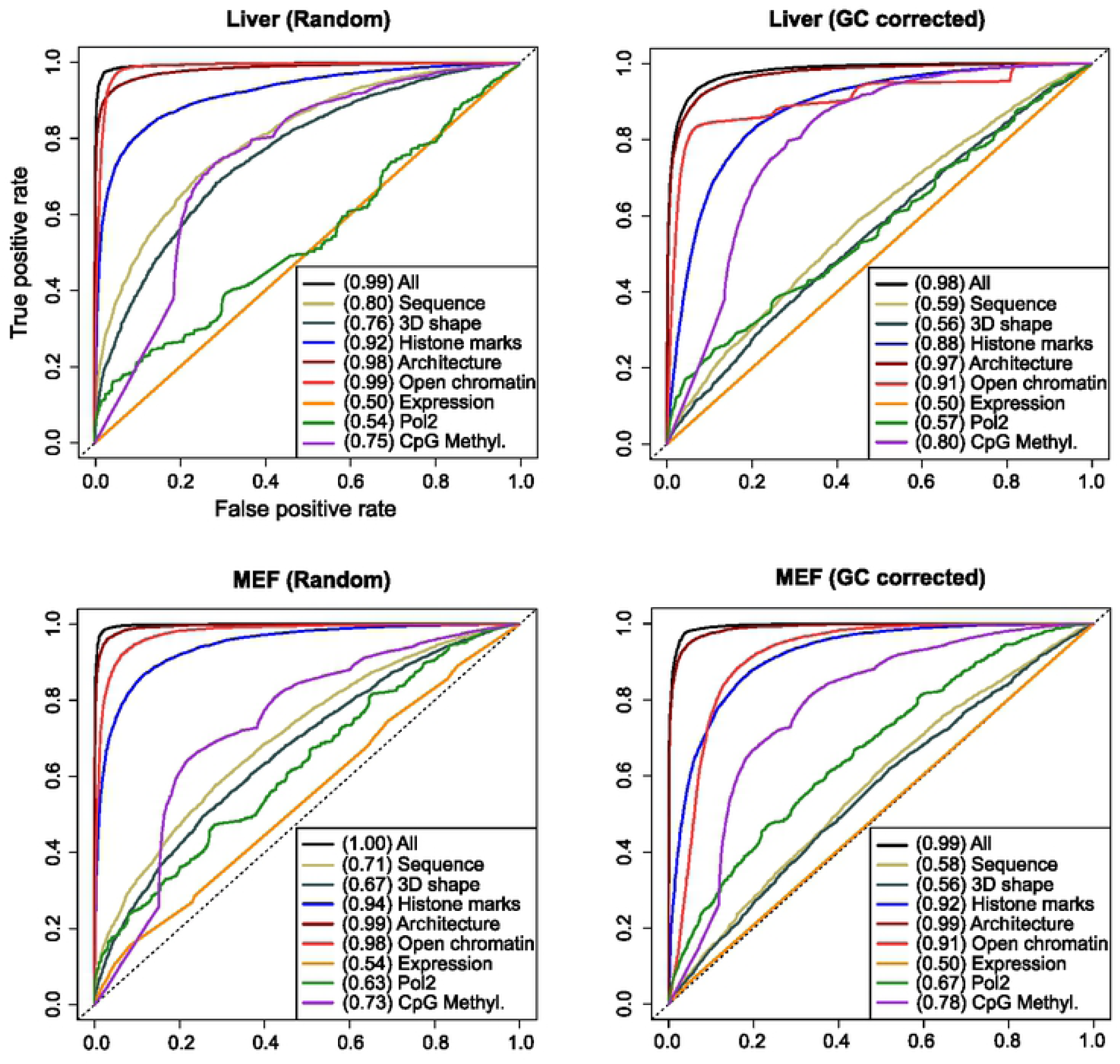
Chromatin features predict TOP2B binding. ROC curves and AUC values for Support Vector Machine models trained on the indicated sets of features (for Naive Bayes models, see S3 Fig)

### Chromatin architecture and open chromatin are the most informative features

To explore whether specific molecular events provide different prediction abilities for TOP2B binding, chromatin features were grouped into 8 categories (Fig 2): open chromatin (DNAse-seq, MNase-seq), histone marks (H3K4me1, H3K4me3, H3K9ac, H3K27ac, H3K27me3, H3K36me3), Pol2 binding (POLR2A ChIP-seq), architectural components (ChIP-seq of CTCF and the cohesin complex components RAD21, STAG1 and STAG2), gene expression (RNA-seq), CpG methylation (WGBS), DNA sequence and DNA shape (measured using DNAshapeR [26]). Then, SVM and NB models were constructed for each of such categories.

Different predictive performances were observed for the grouped molecular events, with similar results across cell types (Tables 1 and 2; Fig 3; S3 Fig). Models trained with architectural components achieved the highest prediction accuracies, ranging from 87.4% to 96.5%, with area under ROC curves (AUC) of 0.96 - 0.99 (Fig 3; S3 Fig). This is in agreement with previous findings of TOP2B associating with the boundaries of TADs [11, 12]. In the same line, models built using open chromatin factors reflect TOP2B preference to bind accessible DNA, showing accuracies of 71.6% - 92.35% and AUC of 0.88 - 0.99 (Tables 1 and 2; Fig 3; S3 Fig). As expected, histone-mark models also obtained reasonably high prediction performances, with accuracies ranging from 78% to 88.6% and AUC of 0.84 - 0.95 (Tables 1 and 2; Fig 3; S3 Fig). The set of histone modifications used to feed these models include a combination of promoter, enhancer and gene body associated marks, which represents common genomic regions where TOP2B tends to localize.

In contrast to the above results, gene expression (measured by RNA-seq) appeared to be the less informative feature for TOP2B binding, with accuracies of around 50% and AUC of 0.5 - 0.54. Interestingly, and despite the observed co-occurrence of TOP2B and Pol2 at gene promoters (Fig effig1; S1 Fig), the model trained with Pol2 binding information also exhibited poor prediction accuracies, ranging from 52.6% to 61.4% and AUC of 0.53 - 0.67. These observations contrast with previous findings of TOP2B being involved in transcriptional regulation [47], and may indicate that other chromatin features, such as histone marks and DNase-seq, are capturing such association.

### The impact of correcting for GC-bias

According to the performances achieved, the effect of training models using GC-corrected background regions is negligible for the categories analyzed. As expected, this is not the case for DNA sequence and DNA shape models, which obtained decreased prediction accuracies when trained with GC-corrected sites as compared to those achieved using pure random regions. While sequence content exhibited a prediction accuracy of 55.4% and 62.9% and AUC of 0.58 - 0.66 for GC-corrected models, these values increased to 64.1% - 74.9% and AUC of 0.71 - 0.84 for random models (Tables 1 and 2; Fig 3; S3 Fig). Similarly, sequence-derived 3D shape models built with GC-corrected regions achieved accuracies from 54% to 57.9% and AUC of 0.56 - 0.6, whereas pure random models obtained accuracies of 61.8% - 73.1% and AUC of 0.67 - 0.78. Given the remarkable resolution of our model in measuring DNA sequence and sequence-derived 3D DNA shape, these results suggest that sequence information alone is only poorly informative of TOP2B binding, and strengthen the idea of topoisomerases acting where DNA topological problems arise.

The predictive performances of models trained with CpG methylation data were also influenced (although only moderately) by the use of GC-corrected sites as background regions. While pure random models achieved modest accuracies, ranging from 64% to 69.5% and AUC of 0.73 - 0.76, GC-corrected models obtained moderately higher accuracies of 69.4% - 75.3% and increased AUC of 0.78 - 0.81 (Tables 1 and 2; Fig 3; S3 Fig). When forcing the G+C content of random regions to match that of TOP2B binding sites, they tend to be located at GC rich sites not occupied by TOP2B. Such regions happen to be hypermethylated as compared with TOP2B sites (S2 Fig), which leads to an increased predictive power of CpG methylation. Given the modest differences found in using random or GC-corrected random regions in model performances, from this point on only GC-corrected random regions were considered.

### Three chromatin features explain TOP2B binding genome wide

With the aim of identifying the most relevant features to explain TOP2B binding, we applied feature selection, which aims at reducing the number of attributes in the data matrix while keeping (or improving) the predictive power of the classifier. We used two feature selection algorithms using 5-fold cross-validation: Fast Correlation Based Filter (FCBF) [33] and Scatter Search (SS) [34, 35] (see Materials and methods). In addition to the accuracy and the AUC values, here we also considered the average number of features selected by each strategy and the stability of each algorithm. The stability (or robustness) evaluates the sensitivity to variations of a feature selection algorithm in the dataset. This issue is especially important in high dimensional domains where strategies may yield to different solutions with similar performance and, therefore, this measure enhances the confidence in the analysis of the results.

Both feature selection algorithms succeeded in finding a subset of features that faithfully explained TOP2B binding in both cell types, with accuracies ranging from 92.1% to 96.11% (Table 3). As a matter of fact, although in mouse liver the two classification approaches (SVM and NB) achieved the same performance for both feature selection algorithms, this was not the case in MEFs, in which the SVM slightly outperformed NB predictions. Regarding the number of selected features, while FCBF needed averages of 46 and 22 features to optimize the prediction of TOP2B in MEF and liver, respectively, SS only needed 3 features in both cell types (S2 Table). It is worth stressing that SS always finds, in every iteration, the same 3 features, achieving a stability score of 1 (Table 3). On the other hand, features selected by FCBF vary from different iterations, which confers a poor robustness score to this strategy.

**Table 3.** Performance of SVM and NB classifiers using the features selected by FCBF and SS strategies.

| Cell type | Algorithm | NB | SVM | #features |
| --- | --- | --- | --- | --- |
| MEF | FCBF | 94.60 ± 0.35 | 96.11 ± 0.28 | 46.80 ± 3.90 |
| | SS | 93.88 ± 0.31 | 95.51 ± 0.33 | 3.00 ± 0.00 |
| Liver | FCBF | 92.62 ± 0.51 | 92.84 ± 1.91 | 22.20 ± 4.38 |
| | SS | 92.90 ± 0.44 | 92.10 ± 1.97 | 3.00 ± 0.00 |

**Table 4.** Performance of cross cell type predictions using models trained with DNase-seq, RAD21 and CTCF. Errors are only shown for models trained and tested with the same cell type due to test data partition.

| Cell type (train/test) | RF | NB | SVM |
| --- | --- | --- | --- |
| MEF | 95.03 ± 0.44 | 93.62 ± 3.76 | 95.22 ± 0.28 |
| MEF/Liver | 91.97 | 89.43 | 92.36 |
| MEF/aB cells | 98.31 | 98.26 | 98.38 |
| Liver | 94.90 ± 0.16 | 91.60 ± 0.51 | 76.15 ± 9.76 |
| Liver/MEF | 89.33 | 89.04 | 93.11 |
| Liver/aB cells | 96.38 | 97.87 | 98.21 |
| aB cells | 98.70 ± 0.22 | 97.63 ± 0.42 | 98.28 ± 0.31 |
| aB cells/MEF | 93.16 | 85.00 | 92.95 |
| aB cells/Liver | 89.08 | 85.49 | 81.19 |

The most selected features for each strategy are shown in Fig 4A. For each histogram and feature, the white bar height indicates the frequency of selection and the black bar height is the Symmetrical Uncertainty (SU) value with respect to the class (TOP2B). SU [48] is a widely used technique that allows to measure the relevance between two variables and can be interpreted as a correlation measure to identify non-linear dependencies between features. Due to the mentioned low stability score obtained with the FCBF approach, only 5 out of the average 46 and 3 out of the average 22 features were selected in every iteration for MEFs and liver, respectively (S2 Table). In MEFs, such 5 features were RAD21, DNase-seq, WGBS, MNase and the DNA shape associated translational parameter Shift. In liver, the three selected features corresponded to DNase-seq, STAG2 and WGBS. It is worth mentioning that the only features selected in 4 of the 5 iterations were the DNA shape associated parameters Slide and Shear in MEFs and Helix Twist in liver. The recurrent selection of such DNA shape features in both cell types may indicate a modest contribution of these parameters to TOP2B binding. In any case, most of the predictive power was sustained by open chromatin features (DNAse-seq, MNase-seq), cohesin complex associated factors (RAD21, STAG2) and to a lesser extent CpG methylation (WGBS).

**Fig 4.**
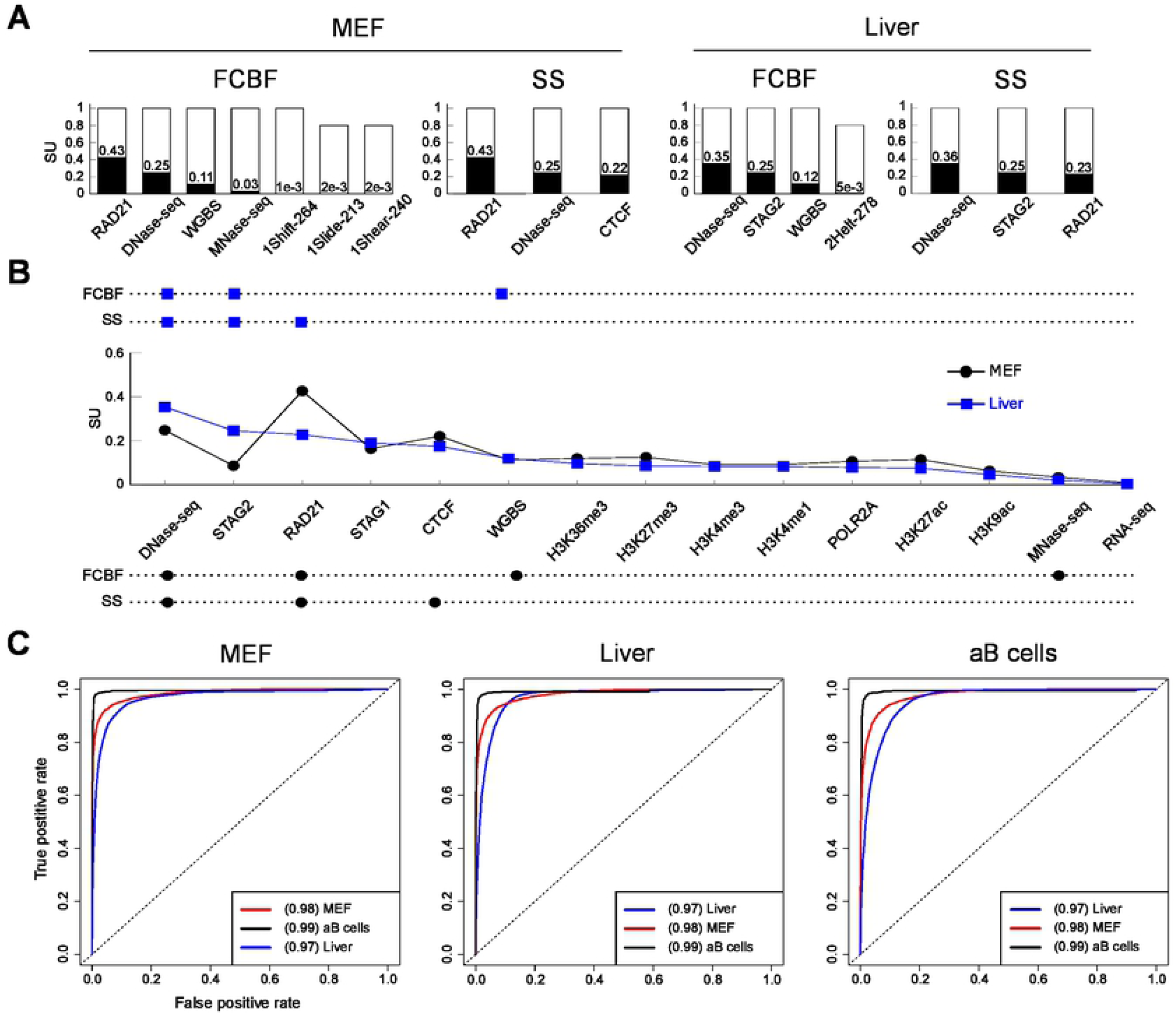
Three features accurately predict TOP2B. **A**. Top features selected by Fast Correlation Based Filter and Scatter Search algorithms. For each histogram and feature, the white bar height indicates the frequency of selection and the black bar height is the Symmetrical Uncertainty (SU) value with respect to the class (TOP2B). Indexes of DNA shape parameters indicate position within the corresponding parameter vector associated to the 300 bp width of modeled TOP2B binding sites (see Materials and methods). **B**. Summary of the most selected features by both algorithms. Top and bottom aligned dots indicate selection of a given feature by the corresponding selection algorithm in liver and MEFs, respectively. In the middle, the SU of each feature is displayed. Only the top fifteen features according to their SU are shown, which happen to match in both cell types. **C**. ROC curves and AUC values for Naive Bayes models trained on either MEF, liver or activated B cells and applied to the three cell types (for Support Vector Machine and Random Forests models, see S4 Fig). Only DNase-seq, RAD21 and CTCF binding data were used for training.

A significantly more robust and consistent selection was achieved when using SS (Fig 4A), where only three features were selected in every iteration for both cell types. Furthermore, DNase-seq and RAD21 were both selected for MEFs and liver, which is in line with above results and highlights the influence of accessible chromatin and cohesin factors for TOP2B binding across cell types. As a matter of fact, while in MEFs SS selected CTCF at every iteration, this was not the case in liver, where the cohesin complex factor STAG2 was always selected (Fig 4A).

A summary of the features selected by FCBS and SS in both cell types is shown in Fig 4B. Top and bottom aligned dots indicate selection of a given feature by the corresponding selection algorithm in liver and MEFs, respectively. In the middle, the relevance (SU) of each feature with respect to TOP2B is displayed. Only the top fifteen features according to their SU are shown, which happen to match in both cell types. With the exception of STAG2, the trends of liver and MEFs are very similar, indicating that the specific contributions of genomic features for TOP2B prediction are conserved across cell types. Furthermore, the results obtained by the SS algorithm strengthen the importance of accessible chromatin and architectural factors for TOP2B binding, and shows that a small number of these features are enough to achieve accurate genome wide predictions.

### Validation in different mouse cell types

We next asked to what extend the three selected features can be used to predict TOP2B binding in cell types that were not used for the training. It is noteworthy to mention that, since data on STAG2 binding is rather scarce, we focused on DNase-seq, RAD21 and CTCF. Although the latter was only selected in MEFs, it was among the most informative features in mouse liver (Fig 4B; S2 Table), and there is a wealth of ChIP-seq data available for this protein. Besides the two cell types considered so far, we included TOP2B binding data from mouse activated B cells (Gene Expression Omnibus GSM2635606). To the best of our knowledge, this dataset is the only publicly available mouse dataset on TOP2B binding for which DNase-seq and ChIP-seq of both CTCF and RAD21 are also accessible (together with the already used from mouse liver and MEFs). We assessed model predictions by training on one cell type and testing prediction accuracy on the rest. Besides SVM and NB, this time we also applied Random Forests (RF) [31], which has been widely applied to genomic data with excellent results [32].

The achieved predictions showed that the cross-cell type applications of the classifiers did not decrease their performances, which acquired accurate predictions independently of the training cell type (Fig 4C; S4 Fig; Table 4). For example, when we applied NB models trained on MEF and mouse liver to activated B cells, we obtained accuracies of 98.3% and 97.9%, respectively, compared to an average of 97.6% using the model trained on activated B cells itself. These results support the idea that TOP2B association with DNase-seq, CTCF and RAD21 can be generalized across cell types.

To further test the above observations, we used genome wide ChIP-seq TOP2B binding in mouse thymocytes obtained in our laboratory [49] and assessed model prediction accuracies on them. This time we generated models using a combination of data from mouse liver, MEFs and activated B cells, which allowed us to better capture the cell type specificities of TOP2B binding in a single classifier. In order to evaluate the specificity of our model, we performed ChIP-seq using two different antibodies (Novus and Santa Cruz). We called peaks using HOMER [22] and identified 49, 974 and 31, 653 TOP2B binding sites for Novus and Santa Cruz, respectively. Then, we applied the model and computed AUC values (Fig 5A). Although modest predictive power was achieved for individual antibodies (AUC of 0.62 and 0.74 for Santa Cruz and Novus, respectively), when applying the model to common peaks of both experiments we obtained quite accurate predictions, with AUC of 0.89.

**Fig 5.**
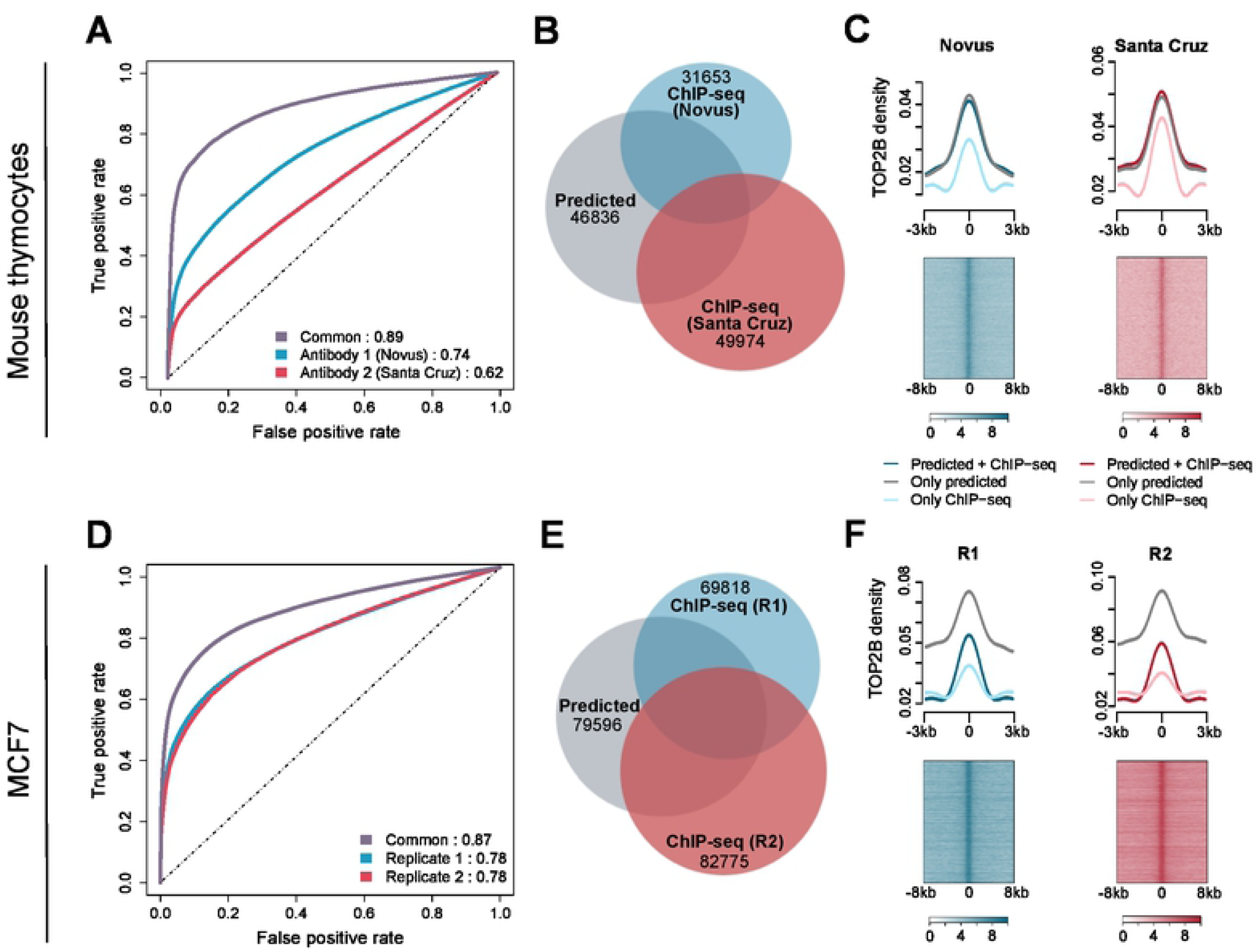
Validation of TOP2B model in mouse thymocytes and human MCF7. **A**. ROC curves and AUC values for the prediction of TOP2B peaks detected using Novus and Santa Cruz antibodies. **B**. Venn diagram showing the overlaps between predicted TOP2B peaks and the two sets of experimental peaks. **C**. Top panel shows average TOP2B ChIP-seq reads enrichment within *±*3 kb of predicted peaks confirmed by HOMER (predicted + ChIP-seq), predicted peaks not detected by HOMER (only predicted) and experimental peaks not predicted by the classifier (Only ChIP-seq). Signals were smoothed using nucleR [39]. Bottom panel shows heatmap representations of TOP2B ChIP-seq reads enrichment within *±*8 kb of predicted peaks not detected by HOMER (only predicted). **D**. ROC curves and AUC values for the prediction of TOP2B peaks in two replicates performed in MCF7. **E**. Venn diagram showing the overlaps between predicted TOP2B peaks and the two sets of experimental peaks. **F**. Top panel shows average TOP2B ChIP-seq reads enrichment within *±*3 kb of predicted peaks confirmed by HOMER (predicted + ChIP-seq), predicted peaks not detected by HOMER (only predicted) and experimental peaks not predicted by the classifier (only ChIP-seq). Bottom panel shows heatmap representations of TOP2B ChIP-seq reads enrichment within *±* 8 kb of predicted peaks not detected by HOMER (only predicted).

**Fig 6.**
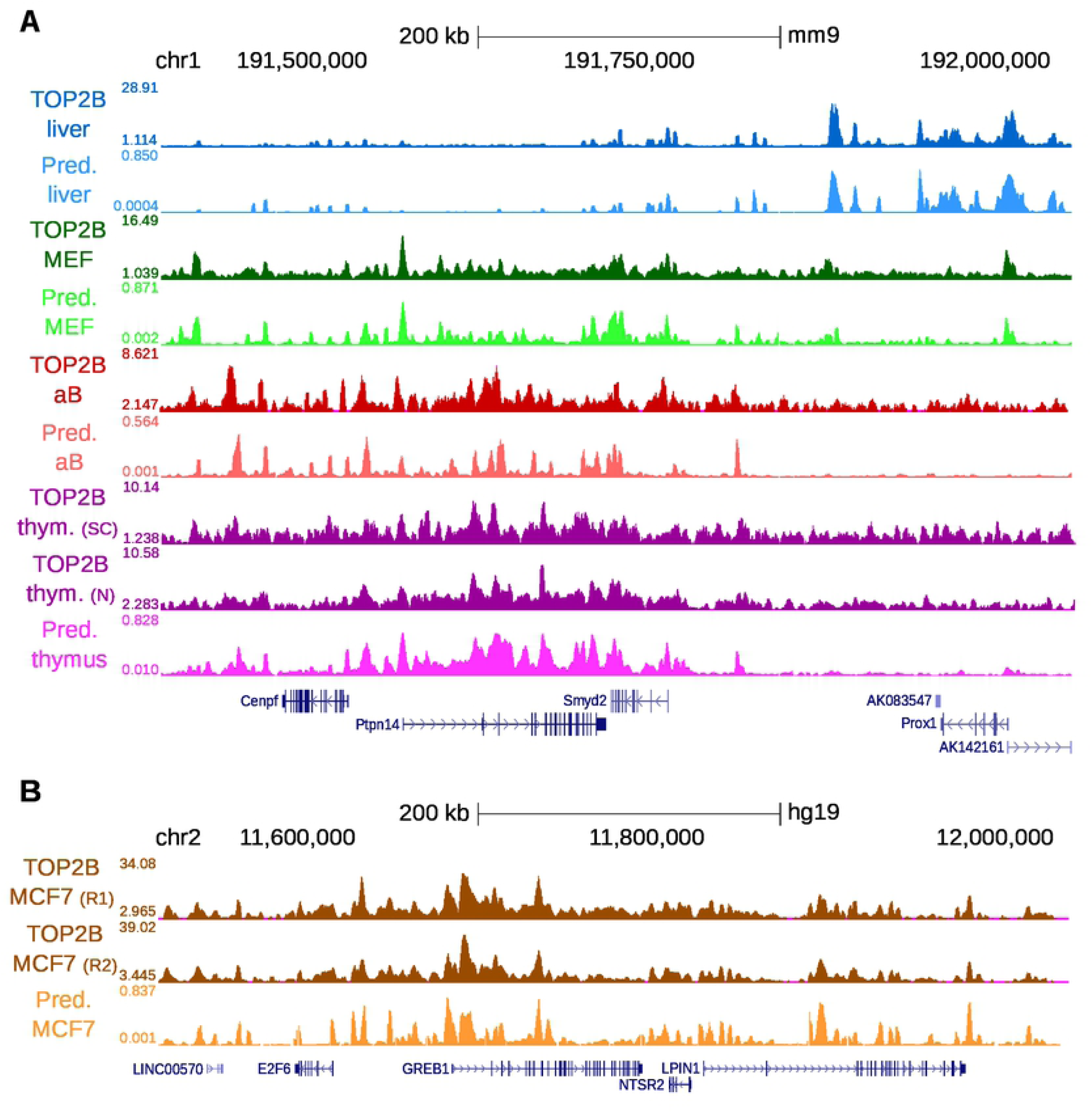
TOP2B predictive and experimental tracks in all cell types used in this study. **A**. Selected region of the mouse genome. Liver, MEFs, activated B cells and thymocytes signals are shown in blue, green, red and purple, respectively. **B**. Selected region of the human genome. MCF7 signals are shown in orange.

Next, we generated genome wide predictions by splitting the genome in 300 bp regions and applying the TOP2B classifier (S5 Fig). We identified 46, 836 regions in the thymus genome with a probability of TOP2B binding above 0.95. Such regions included % of experimental ChIP-seq peaks detected using Novus antibody, 26.7% of those detected using Santa Cruz and 76.2% of common peaks (Fig 5B). In addition, our model detected a set of 24, 566 TOP2B peaks that were not identified by HOMER in any of the two ChIP-seq experiments. In order to assess whether we were potentially detecting false positives, we examined ChIP-seq intensity centered at TOP2B peaks (Fig 5C, top). We displayed TOP2B at three types of peaks: predicted peaks confirmed by HOMER (predicted+ChIP-seq), predicted peaks not detected by HOMER (only predicted) and HOMER peaks not predicted by the classifier (only ChIP-seq). Interestingly, the highest enrichment of TOP2B signal was observed for both types of predicted peaks, independently of them being identified by HOMER or not, indicating that the model faithfully identifies TOP2B binding sites supported by a large amount of ChIP-seq reads, including peaks not called by HOMER. To check if the signal enrichment observed at the latter is an effect of averaging reads around predicted peaks, we represented read enrichment around every region using heatmaps (Fig 5C, bottom). An increased ChIP-seq signal was observed centered at the whole set of predicted peaks not detected by HOMER, suggesting that the model is indeed highly sensitive to TOP2B binding.

### Validation in human cells

In order to test whether our TOP2B binding model can be generalized to other organisms, we performed ChIP-seq of TOP2B in human MCF7 cell line. This time we performed two replicates using the same antibody (Sigma), which allowed us to evaluate to what extend our predictions resemble an experimental replicate. We identified 82, 775 and 69, 818 TOP2B peaks for each replicate, with 37, 774 common peaks. Our model obtained reasonably accurate predictions on both set of peaks, with AUC of 0.78 (Fig 5D). When predicting on the set of common peaks, the model performed considerably better, with AUC of 0.87.

Again, we generated genome wide predictions by tiling the MCF7 genome into 300 bp regions and applying our model to each of them (S5 Fig). We identified 79, 596 regions with a probability of TOP2B binding above 0.95, including 50.7% of experimental peaks from replicate 1, 48.3% of experimental peaks from replicate 2 and 66.7% of common peaks, showing that our predictions behave just as another experimental replicate (Fig 5E). Since our predictions revealed an additional set of 28, 757 peaks that were not detected by HOMER, we assessed whether these corresponded to potential false positives, as described above. We displayed TOP2B ChIP-seq signal at the same three types of peaks: only predicted, only ChIP-seq and both. As expected, predicted peaks confirmed by HOMER showed a high enrichment of TOP2B (Fig 5F, top), indicating that the model is able to identify well-defined binding sites supported by experimental evidence. On the other hand, ChIP-seq peaks not detected by the classifier showed an only modest accumulation of TOP2B. Interestingly, predicted peaks not detected by HOMER showed the highest enrichment of TOP2B, indicating that our model outperforms HOMER in detecting true TOP2B-binding regions. Again, we represented TOP2B ChIP-seq reads using heatmaps in order to test whether the above enrichment is an effect of averaging reads around predicted peaks (Fig 5F, bottom). We observed ChIP-seq signal centered at the whole set of predicted peaks not detected by HOMER, confirming that the model is also sensitive to predict TOP2B binding in human cells.

### TOP2B probability mirrors experimental binding intensity genome wide

Given the remarkable predictive power of our model, we generated virtual tracks genome wide using TOP2B binding probabilities and represented them in a genome browser [37] together with experimental TOP2B ChIP-seq signals. Predictive tracks faithfully simulate ChIP-seq data at the gene level and accurately capture cell type binding differences both in the training and the test sets (Fig 6). It is worth to note that, even though we modeled TOP2B binding using binary classifiers, the similarities between the experimental signals and our predicted probabilities are quite significant. Furthermore, signal comparison with the three chromatin features used as predictors shows that the model is also able to apply learned associations of DNAse-seq, RAD21 and CTCF with TOP2B to precisely generate *bona fide* predictions (S6 and S7 Fig). These results highlight the power of our model and demonstrates that the described approach can be used as a framework for the prediction of genome wide TOP2B signal potentially in any mammalian cell type for which sequencing data on DNAse I hypersensitivity, RAD21 and CTCF are available. Furthermore, this demonstrates that, besides global TOP2B binding, our model can be used to produce virtual probability tracks with which TOP2B accumulation can be analyzed at specific genomic loci.

## Conclusion

TOP2B is a crucial enzyme for DNA metabolism, whose activity is required for a wide range of cellular processes including replication, transcription and genome organization. The few genome-wide maps of TOP2B that are currently available show high colocalization with several chromatin features, but no work has been devoted to comprehensively study predictive features that define TOP2B binding. By applying feature selection techniques, we show that TOP2B binding can be accurately predicted using only three high-throughput sequencing datasets, DNase-seq and ChIP-seq of RAD21 and CTCF, highlighting the importance of TOP2B in the regulation of genome architecture. Since the above datasets are largely available in ENCODE [38] and Roadmap [50], our framework allows the scientific community to predict TOP2B binding in a large set of organisms, cell types, tissues and biological conditions. Finally, we believe that this approach and the generation of virtual probability tracks could be applied to other chromatin binding factors with a similar sequence-independent binding behavior to TOP2B.

## Supporting information

**S1 Fig. Genome browser view for TOP2B ChIP-seq signal and other relevant chromatin features in MEFs**.

**S2 Fig. Enrichment of CpG methylation around TOP2B binding sites in mouse liver and MEFs**. Average of whole genome bisulfite sequencing reads within *±*1 kb of TOP2B binding center, random regions and GC corrected random regions are displayed.

**S3 Fig. Chromatin features predict TOP2B binding**. ROC curves and AUC values for Naive Bayes models trained on the indicated sets of features.

**S4 Fig. ROC curves and AUC values for Support Vector Machine and Random Forests models**. Models were trained on either MEF, liver or activated B cells and applied to the three cell types. Only DNase-seq and ChIP-seq of RAD21 and CTCF were used for training.

**S5 Fig. Framework for the validation of TOP2B model in mouse and human**. First, the genomes of mouse and human were tiled into bins of 300 bp with sliding windows of 50 bp. Then, DNase-seq and ChIP-seq reads of CTCF and RAD21 were scored on those bins (see Materials and methods) and the TOP2B binding model trained on mouse liver, MEFs and activated B cells was applied. Bins having a TOP2B probability higher than 0.95 were classified as TOP2B binding regions. Validation was performed on thymocytes and MCF7 cells for mouse and human, respectively.

**S6 Fig. TOP2B predictive track in mouse liver, activated B cells and MEFs (training cell types)**. From top to bottom, genome browser view for TOP2B ChIP-seq, TOP2B predicted track, CTCF ChIP-seq, RAD21 ChIP-seq and DNase-seq.

**S7 Fig. TOP2B predictive track in human MCF7 and mouse thymocytes (validation cell types)**. From top to bottom, genome browser view for TOP2B ChIP-seq, TOP2B predicted track, CTCF ChIP-seq, RAD21 ChIP-seq and DNase-seq.

**S1 Table. Public and custom sequencing data used in this study**.

**S2 Table. Ranking of features selected by scatter search and fast correlation base filter algorithms**.

## Acknowledgments

We thank A. Herrero-Ruiz and C. Gómez-Marín for valuable comments. All the analyses were performed using custom scripts that were run on the High Perfomance Computing cluster provided by the Centro Informático Científico de Andalucía (CICA).

## References

1. Vos SM, Tretter EM, Schmidt BH, Berger JM. All tangled up: How cells direct, manage and exploit topoisomerase function; 2011.

2. Nitiss JL. DNA topoisomerase II and its growing repertoire of biological functions. Nature Reviews Cancer. 2009;9(5):327–337. doi:10.1038/nrc2608.

3. Pommier Y, Sun Y, Huang SyN, Nitiss JL. Roles of eukaryotic topoisomerases in transcription, replication and genomic stability. Nature Reviews Molecular Cell Biology. 2016;17(11):703–721. doi:10.1038/nrm.2016.111.

4. Deweese JE, Osheroff N. The DNA cleavage reaction of topoisomerase II: Wolf in sheep’s clothing. Nucleic Acids Research. 2009;37(3):738–748. doi:10.1093/nar/gkn937.

5. Akimitsu N, Kamura K, Toné S, Sakaguchi A, Kikuchi A, Hamamoto H, et al. Induction of apoptosis by depletion of DNA topoisomerase II*α* in mammalian cells. Biochemical and Biophysical Research Communications. 2003;307(2):301–307. doi:10.1016/S0006-291X(03)01169-0.

6. Thakurela S, Garding A, Jung J, Schubeler D, Burger L, Tiwari VK. Gene regulation and priming by topoisomerase IIalpha in embryonic stem cells. Nat Commun. 2013;4:2478. doi:10.1038/ncomms3478.

7. Dereuddre S, Delaporte C, Jacquemin-Sablon A. Role of topoisomerase II*β* in the resistance of 9-OH-ellipticine-resistant Chinese hamster fibroblasts to topoisomerase II inhibitors. Cancer Research. 1997;57(19):4301–4308.

8. Capranico G, Tinelli S, Austin CA, Fisher ML, Zunino F. Different patterns of gene expression of topoisomerase II isoforms in differentiated tissues during murine development. BBA - Gene Structure and Expression. 1992;1132(1):43–48. doi:10.1016/0167-4781(92)90050-A.

9. Madabhushi R, Gao F, Pfenning AR, Pan L, Yamakawa S, Seo J, et al. Activity-Induced DNA Breaks Govern the Expression of Neuronal Early-Response Genes. Cell. 2015;161(7):1592–1605. doi:10.1016/j.cell.2015.05.032.

10. Manville CM, Smith K, Sondka Z, Rance H, Cockell S, Cowell IG, et al. Genome-wide ChIP-seq analysis of human TOP2B occupancy in MCF7 breast cancer epithelial cells. Biology Open. 2015;4(11):1436–1447. doi:10.1242/bio.014308.

11. Uusküla-Reimand L, Hou H, Samavarchi-Tehrani P, Rudan MV, Liang M, Medina-Rivera A, et al. Topoisomerase II beta interacts with cohesin and CTCF at topological domain borders. Genome Biol. 2016;17(1):1–22. doi:10.1186/s13059-016-1043-8.

12. Canela A, Maman Y, Jung S, Wong N, Callen E, Day A, et al. Genome Organization Drives Chromosome Fragility. Cell. 2017;170(3):507–521.e18. doi:10.1016/j.cell.2017.06.034.

13. Dellino GI, Palluzzi F, Chiariello AM, Piccioni R, Bianco S, Furia L, et al. Release of paused RNA polymerase II at specific loci favors DNA double-strand-break formation and promotes cancer translocations. Nature Genetics. 2019;doi:10.1038/s41588-019-0421-z.

14. Ciccia A, Elledge SJ. The DNA Damage Response: Making It Safe to Play with Knives; 2010.

15. Polo SE, Jackson SP. Dynamics of DNA damage response proteins at DNA breaks: A focus on protein modifications; 2011.

16. Chapman JR, Taylor MRG, Boulton SJ. Playing the End Game: DNA Double-Strand Break Repair Pathway Choice; 2012.

17. Canela A, Maman Y, Huang SyN, Wutz G, Tang W, Zagnoli-Vieira G, et al. Topoisomerase II-Induced Chromosome Breakage and Translocation Is Determined by Chromosome Architecture and Transcriptional Activity. Molecular Cell. 2019;doi:10.1016/j.molcel.2019.04.030.

18. Gothe HJ, Bouwman BAM, Gusmao EG, Piccinno R, Petrosino G, Sayols S, et al. Spatial Chromosome Folding and Active Transcription Drive DNA Fragility and Formation of Oncogenic MLL Translocations. Molecular Cell. 2019;doi:10.1016/j.molcel.2019.05.015.

19. Langmead B, Trapnell C, Pop M, Salzberg S. Ultrafast and memory-efficient alignment of short DNA sequences to the human genome. Genome biology. 2009;10(3):R25. doi:10.1186/gb-2009-10-3-r25.

20. Hahne F, Ivanek R. Visualizing genomic data using Gviz and bioconductor. In: Methods in Molecular Biology; 2016.

21. Ou J, Zhu LJ. trackViewer: a Bioconductor package for interactive and integrative visualization of multi-omics data; 2019.

22. Heinz S, Benner C, Spann N, Bertolino E, Lin YC, Laslo P, et al. Simple Combinations of Lineage-Determining Transcription Factors Prime cis-Regulatory Elements Required for Macrophage and B Cell Identities. Molecular Cell. 2010;doi:10.1016/j.molcel.2010.05.004.

23. Amemiya HM, Kundaje A, Boyle AP. The ENCODE Blacklist: Identification of Problematic Regions of the Genome. Scientific Reports. 2019;doi:10.1038/s41598-019-45839-z.

24. Comoglio F, Paro R. Combinatorial Modeling of Chromatin Features Quantitatively Predicts DNA Replication Timing in Drosophila. PLoS Computational Biology. 2014;10(1). doi:10.1371/journal.pcbi.1003419.

25. Lister R, Mukamel EA, Nery JR, Urich M, Puddifoot CA, Johnson ND, et al. Global epigenomic reconfiguration during mammalian brain development. Science. 2013;doi:10.1126/science.1237905.

26. Chiu TP, Comoglio F, Zhou T, Yang L, Paro R, Rohs R. DNAshapeR: An R/Bioconductor package for DNA shape prediction and feature encoding. Bioinformatics. 2016;doi:10.1093/bioinformatics/btv735.

27. Li J, Sagendorf JM, Chiu TP, Pasi M, Perez A, Rohs R. Expanding the repertoire of DNA shape features for genome-scale studies of transcription factor binding. Nucleic Acids Research. 2017;doi:10.1093/nar/gkx1145.

28. Zhou T, Shen N, Yang L, Abe N, Horton J, Mann RS, et al. Quantitative modeling of transcription factor binding specificities using DNA shape. Proceedings of the National Academy of Sciences. 2015;112(15):4654–4659. doi:10.1073/pnas.1422023112.

29. Duda RO, Hart PE. Pattern classification and scene analysis. New York, USA: John Wiley & Sons; 1973.

30. Vapnik VN. Statistical Learning Theory. Wiley-Interscience; 1998.

31. Breiman L. Random Forests. Machine Learning. 2001;45:5–32.

32. Chen X, Ishwaran H. Random forests for genomic data analysis; 2012.

33. Yu L, Liu H. Efficient Feature Selection via Analysis of Relevance and Redundancy. Journal of Machine Learning Research. 2004;5:1205–1224.

34. García López FC, García Torres M, Moreno Pérez JA, Moreno Vega JM. Scatter Search for the Feature Selection Problem. In: Conejo R, Urretavizcaya M, Pérez-de-la Cruz JL, editors. Current Topics in Artificial Intelligence. Berlin, Heidelberg: Springer Berlin Heidelberg; 2004. p. 517–525.

35. García-López FC, García-Torres M, Melián-Batista B, Pérez JAM, Moreno-Vega JM. Solving the Feature Selection Problem by a Parallel Scatter Search. European Journal of Operations Research. 2006;169(2):477–489.

36. Hall MA. Correlation-based Feature Subset Selection for Machine Learning. University of Waikato; 1998.

37. Kent WJ, Sugnet CW, Furey TS, Roskin KM, Pringle TH, Zahler AM, et al. The Human Genome Browser at UCSC. Genome Research. 2002;doi:10.1101/gr.229102.

38. Consortium EP. An integrated encyclopedia of DNA elements in the human genome. Nature. 2012;489(7414):57–74. doi:10.1038/nature11247.

39. Flores O, Orozco M. nucleR: A package for non-parametric nucleosome positioning. Bioinformatics. 2011;doi:10.1093/bioinformatics/btr345.

40. Kim SH, Ganji M, Kim E, van der Torre J, Abbondanzieri E, Dekker C. DNA sequence encodes the position of DNA supercoils. eLife. 2018;doi:10.7554/eLife.36557.

41. Mathelier A, Xin B, Chiu TP, Yang L, Rohs R, Wasserman WW. DNA Shape Features Improve Transcription Factor Binding Site Predictions In Vivo. Cell Systems. 2016;3(3):278–286.e4. doi:10.1016/j.cels.2016.07.001.

42. Rao S, Chiu TP, Kribelbauer JF, Mann RS, Bussemaker HJ, Rohs R. Systematic prediction of DNA shape changes due to CpG methylation explains epigenetic effects on protein-DNA binding. Epigenetics and Chromatin. 2018;doi:10.1186/s13072-018-0174-4.

43. Jones PA. Functions of DNA methylation: islands, start sites, gene bodies and beyond. Nature Reviews Genetics. 2012;13(7):484–492. doi:10.1038/nrg3230.

44. Vinson C, Chatterjee R. CG methylation. Epigenomics. 2012;4(6):655–663. doi:10.2217/epi.12.55.

45. Rube HT, Lee W, Hejna M, Chen H, Yasui DH, Hess JF, et al. Sequence features accurately predict genome-wide MeCP2 binding in vivo. Nature Communications. 2016;doi:10.1038/ncomms11025.

46. Zhou T, Yang L, Lu Y, Dror I, Dantas Machado AC, Ghane T, et al. DNAshape: a method for the high-throughput prediction of DNA structural features on a genomic scale. Nucleic acids research. 2013;41(Web Server issue). doi:10.1093/nar/gkt437.

47. Tiwari VK, Burger L, Nikoletopoulou V, Deogracias R, Thakurela S, Wirbelauer C, et al. Target genes of Topoisomerase II regulate neuronal survival and are defined by their chromatin state. Proceedings of the National Academy of Sciences. 2012;109(16):E934–E943. doi:10.1073/pnas.1119798109.

48. Witten IH, Frank E, Hall MA, Pal CJ. Data Mining: Practical Machine Learning Tools and Techniques; 2016.

49. Álvarez-Quilón A, Terrón-Bautista J, Delgado-Sainz I, Serrano-Benítez A, Romero-Granados R, Martínez-García PM, et al. Endogenous topoisomerase II-mediated DNA breaks drive thymic cancer predisposition linked to ATM deficiency. Nature Communications. 2020;doi:10.1038/s41467-020-14638-w.

50. Roadmap Epigenomics Consortium. Integrative analysis of 111 reference human epigenomes. Nature. 2015;doi:10.1038/nature14248.

